# A comprehensive map of missense trafficking variants in rhodopsin and their response to pharmacologic correction

**DOI:** 10.1101/2025.02.27.640335

**Authors:** Kannan V. Manian, Connor H. Ludwig, Yan Zhao, Nathan Abell, Xiaoping Yang, David E. Root, Matthew L. Albert, Jason Comander

## Abstract

Rhodopsin (*RHO*) missense variants are a leading cause of autosomal dominant retinitis pigmentosa (adRP), a progressive retinal degeneration with no currently approved therapies. Interpreting the pathogenicity of the growing number of identified *RHO* variants is a major clinical challenge, and understanding their disease mechanisms is essential for developing effective therapies. Here, we present a high-resolution map of *RHO* missense variant trafficking using two complementary deep mutational scanning (DMS) approaches based on a surface abundance immunoassay and a membrane proximity assay. We generated a comprehensive dataset encompassing all 6,612 possible single-residue missense variants, revealing a strong correlation between the two methods. Over 700 variants were identified with pathogenic trafficking scores, significantly expanding the number of *RHO* variants with functional evidence supporting pathogenicity. We demonstrate a high concordance between the trafficking scores and ClinVar pathogenicity classifications, highlighting this approach’s utility in resolving variants of uncertain significance (VUS). The data also identified structurally clustered trafficking-deficient variants, predominantly within the N-terminal region and second extracellular loop, in and above the extracellular/intradiscal beta-plug region. Furthermore, we evaluated the efficacy of the non-retinoid pharmacological chaperone YC-001, observing significant rescue of trafficking defects in a majority of mistrafficking variants. This comprehensive functional map of *RHO* missense variants provides a valuable resource for pathogenicity assessment, genotype-phenotype correlations, and the development of targeted therapeutic strategies for *RHO*-adRP, paving the way for improved diagnosis and treatment for patients.

## Introduction

The rhodopsin (*RHO*) gene holds a unique place in structural biology and genetics. As the first G-protein-coupled receptor (GPCR) to be crystallized^1^, it provided a foundational model for GPCR structure-function relationships, influencing drug discovery across diverse therapeutic areas^2,3^. It was also among the first human disease genes identified and, as a GPCR essential for rod photoreceptor function^4^, the first gene identified that causes an inherited retinal degeneration disease^5,6^. More than three decades of research, spanning mutagenesis studies^7^, biophysical characterization^8^, and high-resolution structural analyses^9^, have shaped our understanding of rhodopsin’s function and its role in the progressive, degenerative retinal disease autosomal dominant retinitis pigmentosa (*RHO*-adRP)^10,11^. Despite these advances, a comprehensive map of *RHO* missense variants and their functional consequences remains elusive. Deep mutational scanning (DMS) has emerged as a transformative approach for systematically mapping structure-function relationships and measuring disease-associated phenotypes across entire mutational landscapes. This approach has been successfully applied to proteins with well-defined folding and trafficking pathways, including ion channels^12,13^, tumor suppressors^14,15^, and metabolic enzymes^16,17^, to resolve variant pathogenicity and identify therapeutic entry points. Given rhodopsin’s central place in vision research, the genetic complexity of *RHO*-adRP, and the emergence of clinical trials for *RHO*-adRP patients, a systematic characterization of *RHO* variants is increasingly needed.

Pathogenic *RHO* variants that cause *RHO*-adRP span multiple mechanistic classes, with misfolding and endoplasmic reticulum (ER) retention representing the most prevalent pathogenic mechanism. This class of variants disrupts rhodopsin’s trafficking through the secretory pathway, leading to ER stress and photoreceptor cell death^10,18^. While some functional data exist for well-characterized variants, most *RHO* variants remain unclassified, limiting their utility for genetic counseling, prognostic assessment, and targeted therapeutic development. Clinical genetic testing has expanded the identification of *RHO* variants, yet many remain variants of uncertain significance (VUS), creating barriers to precision medicine^19^. The American College of Medical Genetics and Genomics (ACMG) scoring system allows for well-validated functional assays to contribute to variant classification^20–23^. Prior studies have successfully used heterologous cells to demonstrate cell trafficking defects of *RHO* variants, including a set of 210 literature and clinical *RHO* variants^24^, a set of 123 pathogenic *RHO* variants^25^, and 700 transmembrane domain *RHO* variants^26^. However, no systematic effort has comprehensively resolved the functional consequences of all *RHO* missense variants. A high-resolution functional landscape of *RHO* variants would enable pathogenicity assessment, inform genotype-phenotype correlations, and provide a foundation for novel therapeutic strategies.

Herein, we apply two complementary DMS approaches to systematically profile the trafficking properties of all possible *RHO* missense variants, generating a comprehensive functional map of *RHO* mistrafficking variants. We identify structurally clustered, trafficking-deficient variants enriched in pathogenic annotations and demonstrate that a previously reported small-molecule corrector YC-001^27^ can rescue the majority of these variants. These data enhance our understanding of rhodopsin biology, provide a scalable framework for variant classification, and lay the foundation for precision ophthalmology approaches to *RHO*-adRP. By integrating decades of research with modern high-throughput functional genomics, we aim to bridge the gap between genetic discovery and therapeutic intervention, expanding opportunities for individualized treatment strategies in inherited retinal degeneration diseases.

## Results

### Dual deep mutational scanning approaches for RHO surface trafficking

To systematically identify single-residue *RHO* variants with folding and trafficking defects, we developed two complementary, highly scalable, multiplex, HEK293T-cell based assays capable of measuring RHO surface trafficking. The first method, henceforth ‘Method 1’, used a flow cytometry-based immunoassay to quantify trafficking and steady-state surface abundance of RHO (**Figure 1A**). Briefly, cells were transduced with a lentiviral library (**Figure S1A**) containing all single-residue missense, nonsense, and synonymous *RHO* variants (constitutively expressed), using a low multiplicity of infection to ensure integration of only one variant per cell. Approximately 500 cells per variant were transduced to buffer against cell-to-cell variability in the lentiviral integration site, and successfully transduced cells were selected with puromycin. For the assay, cells were fixed and stained with an antibody recognizing the first 10 residues of the extracellular N-terminus of RHO. Cells were then subjected to fluorescence-activated cell sorting (FACS) for separation into high- and low-fluorescence bins (Methods). Genomic DNA from each bin was purified, and *RHO* variants were read out by next-generation sequencing (NGS) and quantified.

**Figure 1.**
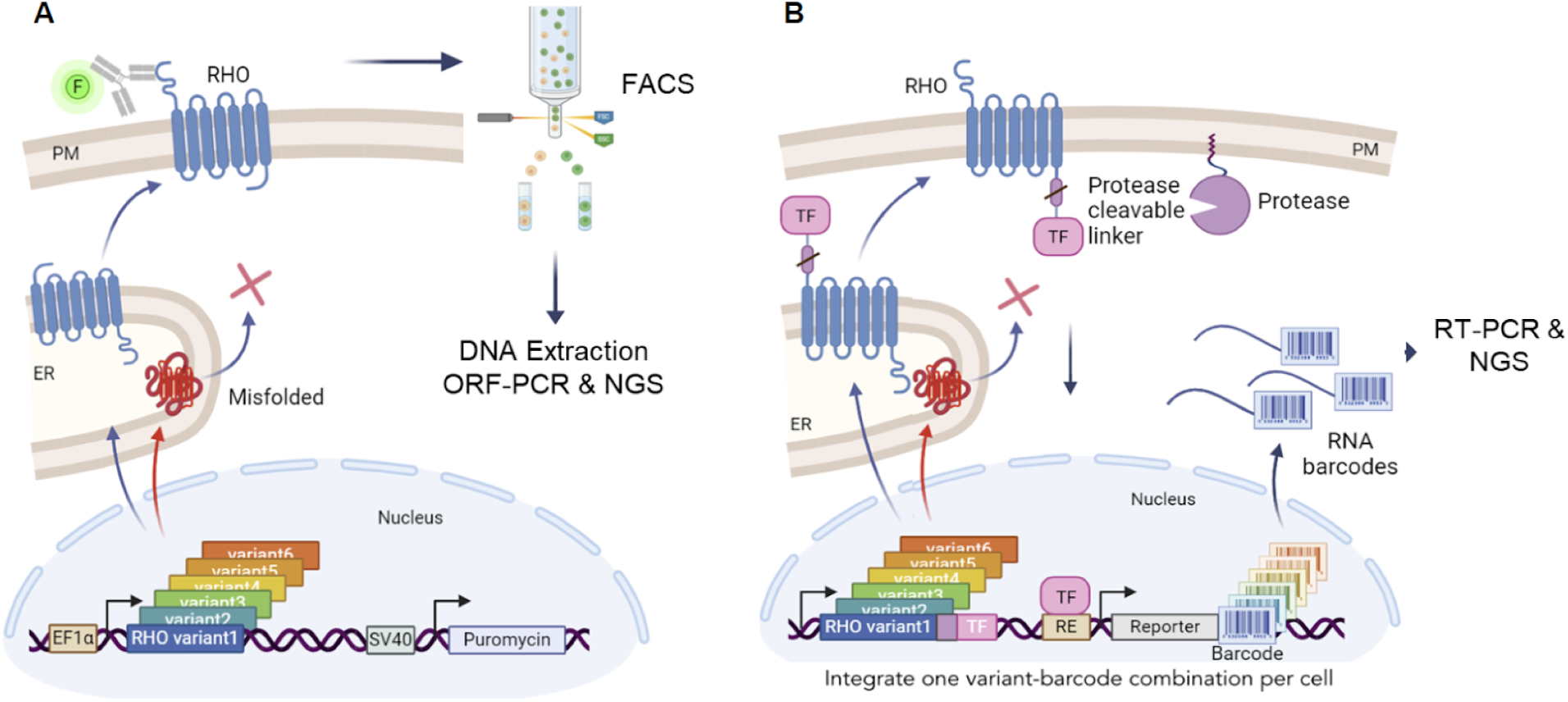
Two deep mutational scanning methods to measure RHO variant surface trafficking. (**A**) In Method 1, a single-site saturation mutagenesis library of *RHO* is transduced into HEK293T cells. *RHO* variants are constitutively expressed in their native, untagged form and selected using puromycin. Cells are fixed, immunostained with a fluorophore-conjugated antibody that recognizes the extracellular N-terminus of RHO, and subjected to fluorescence-activated cell sorting (FACS) to separate and sort the top 20% RHO^High^ expression bin and the bottom 20% RHO^Low^ expression bin. Genomic DNA from each population is reverse-crosslinked, *RHO* variant sequences are amplified with open reading frame polymerase chain reaction (ORF-PCR), and variant frequencies in each population are quantified using next generation sequencing (NGS). (**B**) In Method 2, a single-site saturation mutagenesis library of *RHO* is integrated at single copy into a specific landing pad in the H11 safe harbor locus of HEK293T cells. *RHO* variants are inducibly expressed using the Tet-ON system and are expressed with a C-terminally fused protease-cleavable linker followed by a transcription factor (TF). Proper folding of RHO in the endoplasmic reticulum (ER) and trafficking to the plasma membrane (PM) positions it proximal to a membrane-anchored protease that releases the TF to translocate into the nucleus, thus allowing binding to a specific DNA response element (RE) and initiation of transcription of a variant-specific barcode. RNA barcodes are purified from cell lysates and used in a reverse transcription reaction to make complementary DNA barcodes, which are amplified by PCR and read out by NGS (not shown).

The second method, henceforth ‘Method 2’, used a transcriptional reporter assay to measure surface trafficking of RHO (**Figure 1B**). Cells harboring a landing pad in the *H11* locus were co-transfected with a plasmid library containing all single-residue *RHO* missense and nonsense variants as well as a plasmid encoding the Bxb1 integrase. This integrase ensured landing pad-specific, single-copy integration of an all-in-one library construct (**Figure S1B**) containing the following elements: (i) a constitutively expressed plasma membrane-anchored protease, (ii) one inducibly (Tet-On) expressed *RHO* variant fused via a protease-cleavable linker to a transcription factor, and (iii) a reporter gene that produced a unique RNA barcode upon transcription factor binding that served as an identifier for the encoded genetic variant. Proper trafficking of RHO to the plasma membrane resulted in protease-mediated release of the transcription factor, which once free translocated to the nucleus and activated transcription of the DNA-encoded RNA barcode. RNA was harvested from cells and converted to cDNA, and the transcribed barcodes were read out by NGS.

For each method, sequencing counts were modeled using a negative binomial generalized linear mixed-effects model (NBGLMM) to estimate variant effects. Effect sizes describing the log_2_ fold change between variant and wild-type were adjusted to a zero-to-one log scale, where zero represents the global mean of nonsense variant surface trafficking scores and one represents wild-type or synonymous variant scores.

### Correlation of variant effects using complementary DMS approaches

For Method 1, we performed a single library transduction and two replicate FACS assays. We recovered 6,572 of the 6,612 missense variants (99.4%), 346 of the 348 nonsense variants, and 328 of the 348 synonymous variants. For Method 2, we performed a single library transfection and eight replicate assays. We recovered 6,611 of the 6,612 missense variants (>99.9%) and all 348 of the nonsense variants, with a median of 57.5 unique barcodes per variant per RNA-seq sample. For both methods, surface trafficking data were bimodally distributed (**Figure 2A**), with 1,533 (23.3%) and 1,490 (22.5%) missense variants exhibiting significantly lower surface trafficking than wild-type in Method 1 and Method 2, respectively, based on a 95% confidence interval.

**Figure 2.**
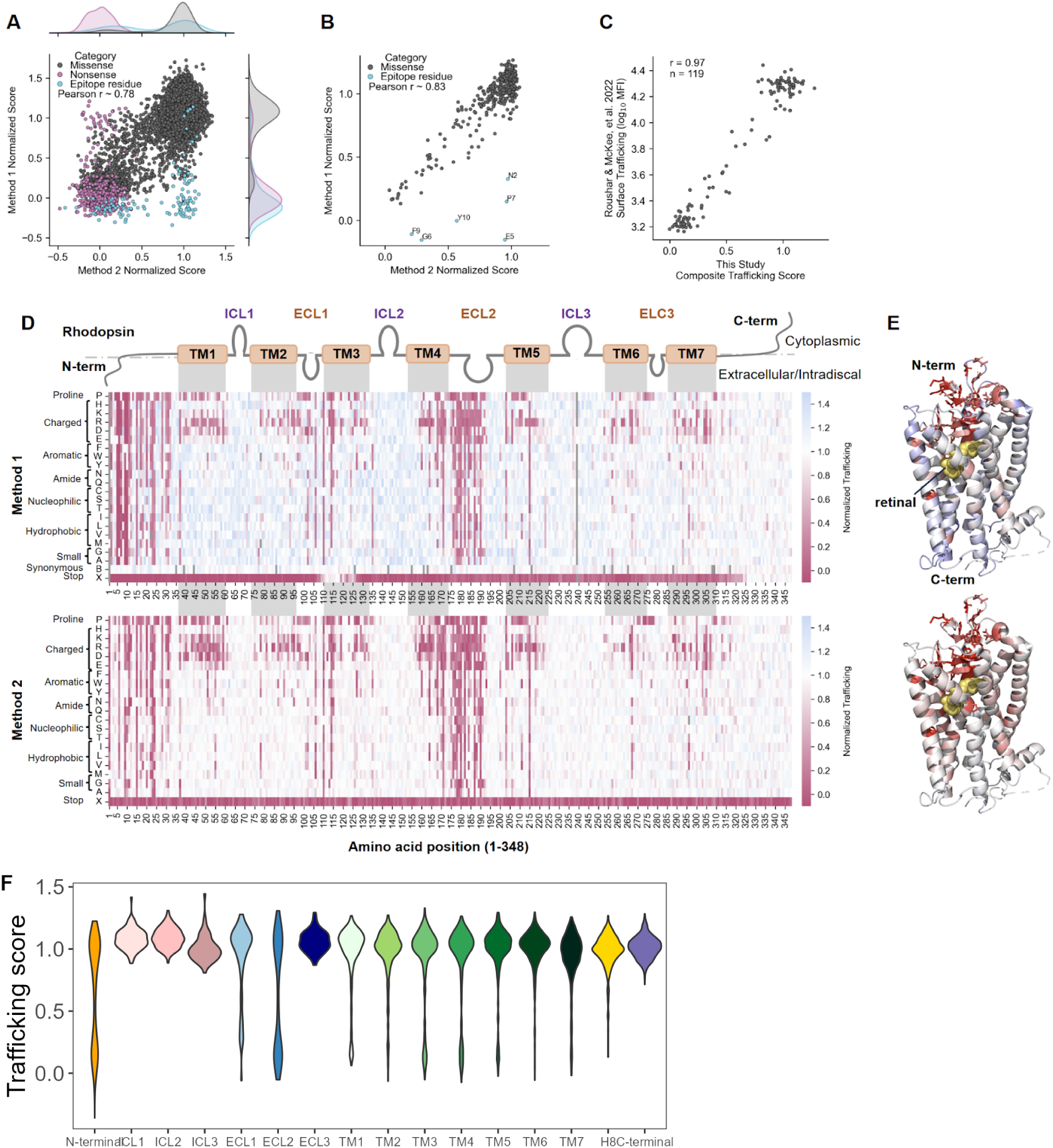
Extracellular-facing residues positioned above the retinal-binding pocket are particularly mutationally intolerant. (**A**) Scatterplot showing strong agreement between deep mutational scanning (DMS) scores for Method 1 (y-axis) and Method 2 (x-axis) at the variant level, with adjunct kernel density estimations showing the distribution of scores for each dataset. Variants affected by Method 1- and Method 2-specific artifacts are highlighted in blue and pink, respectively (Pearson’s r = 0.78). (**B**) Scatterplot showing strong agreement between average DMS scores for Method 1 (y-axis) and Method 2 (x-axis) at the position level. Nonsense variants (affected by Method 2-specific artifacts) are excluded, and variants affected by Method 1-specific artifacts are highlighted in blue (Pearson’s r = 0.83). (**C**) Scatterplot showing strong agreement between the composite trafficking scores from this study (x-axis) and the surface trafficking measurements (119 missense variants) from a flow cytometry-based HEK293T immunoassay as presented in Roushar & McKee, et al. 2022 (y-axis) (r = 0.97)^25^. (**D**) Heatmaps showing surface trafficking for Method 1 (top) and Method 2 (bottom) for all missense and nonsense variants. The x-axis indicates the position along the RHO protein, with domain boundaries indicated (TM = transmembrane, ICL = intracellular loop, ECL = extracellular loop). The y-axis shows the amino acid substituted at each position. RHO surface trafficking is normalized on a log scale from 0 (mean of non-trafficking, nonsense variant negative controls) to 1 (wild-type RHO surface trafficking, positive control) and is indicated by cell color, with white representing trafficking similar to wild-type, pink representing a trafficking defect relative to wild-type, and blue representing a trafficking increase relative to wild-type. Missing measurements are indicated in gray. (**E**) RHO structure (PDB: 1F88) colored according to the average DMS score at each position (excluding nonsense variants) for Method 1 (top) and Method 2 (bottom). (**F**) Distributions of composite trafficking scores (from meta-analysis, see Methods) for each domain of RHO.

The use of two complementary DMS approaches provided an opportunity to assess RHO trafficking with independent methodologies. Despite the methodological distinctions between Method 1 and Method 2 (**Figure 1**), we observed strong agreement between Method 1 and Method 2 trafficking scores on a per-variant level (**Figure 2A**, Pearson r = 0.78, p < 0.01) and on a per-residue level (**Figure 2B**, Pearson r = 0.83, p < 0.01). Moreover, the use of complementary methodologies enabled us to fill gaps where technical limitations affected one assay but not the other. Notably, the most discordant variants for Method 1 were within the N-terminal, surface-exposed epitope region (residues 2-10), which interfered with antibody binding and accounted for the observed low signal (**Figure 2A-B, blue**). By contrast, the use of a transcription factor fused to the C-terminus of RHO in Method 2 accounted for the inability to detect truncation variants that were nonetheless capable of trafficking to the cell surface (**Figure 2A, pink**). Method 1 produced synonymous variant trafficking scores at every position in the library, and they demonstrated a strong separation from the scores of nonsense variants. While the vast majority of nonsense variants failed to traffic to the plasma membrane, Method 1 revealed a subset of nonsense variants from residues 110-120 (after the first two transmembrane domains) and 320-348 (unstructured cytosolic C-terminus) that had near-normal surface trafficking. Consistent with the pooled assay results, immunofluorescence and flow cytometry analysis of individual variants confirmed the moderate surface trafficking for nonsense variants at G114 and F116 variants but showed no detectable surface trafficking for that at W126 (**Figure S2**).

We then therefore performed a meta-analysis using the Method 1 and Method 2 trafficking scores in order to obtain a pooled, more precise trafficking metric for each variant. The variants with the top 5% most discordant scores were flagged (Methods, I^2^ heterogeneity threshold of 93). Our composite trafficking scores strongly correlated with those measured by a flow cytometry-based HEK293T immunoassay (similar to Method 1) in a recent study of 119 pathogenic *RHO* missense variants^25^ (**Figure 2C**, Pearson r = 0.97, p < 0.01).

### Spatial grouping of variant effects

Variants with surface trafficking defects were not evenly distributed across the RHO protein. Strikingly, a large fraction of substitutions within the N-terminal region and second extracellular loop (ECL2) produced severe trafficking defects (**Figure 2D-F**). These regions cover the retinal-binding pocket, harbor crucial residues for proper protein folding and glycosylation, and coordinate key interactions with ECL1 and ECL3^28^. For instance, all substitutions were deleterious at C110 and C187, which form a critical disulfide bridge between ECL1 and ECL2 ^29^. Additionally, the seven transmembrane structure of RHO is visible in the heatmap, with intolerance to transmembrane (TM) domain substitutions to proline, a canonical helix-breaker, and to charged amino acids, which are electrostatically disfavored in the hydrophobic lipid bilayer (**Figure 2D-E**), as has been observed for other GPCRs^30–33^. The cytoplasmic C-terminus of RHO is critical for interactions responsible for ciliary and rod photoreceptor outer segment targeting, and this compartment is not available in HEK293T cells. Accordingly, both DMS methods showed normal trafficking of missense and nonsense variants within the RHO C-terminal tail (326-348) (**Figure 2D-F**); additional assays will be needed to detect pathogenic variants in this region.

### Concordance with mechanistic class labels

Mechanistic class labels for *RHO* variants have developed over time, and one system defines seven mechanistic classes for pathogenic *RHO* variants based on their biochemical and cellular dysfunction: altered post golgi trafficking and outer segment targeting (Class 1), misfolding with ER retention and instability (2), disrupted vesicular trafficking and endocytosis (3), altered post-translational modifications and stability (4), altered transducin activation (5), constitutive activation (6), and dimerization deficiency (7)^10,24^. Most pathogenic variants that have been assigned a mechanistic class label misfold and are retained in the ER (Class 2), which the assays used in this study are designed to detect as a defect in surface trafficking. To assess the ability of both methods to identify surface trafficking defects for Class 2 variants, we overlaid biochemical mechanistic class labels for the small set of 91 literature-annotated variants^10,24^. As expected, annotated variants with the greatest surface trafficking defects belonged to Class 2, including the founder variant P23H (**Figure 3A-B**). While most annotated Class 2 variants mistrafficked, a few had no trafficking defects, suggesting consideration for reclassification. We observed a range of trafficking scores for Class 3 variants, which have only been described at R135 (**Figure 3A**). R135W, the Class 3 variant with the most severe trafficking defect, has been described as misfolded and ER-retained, which are hallmarks of Class 2^34^ (**Figure 3A**). Misfolding and ER retention have not previously been observed for R135L and R135P, which had some of the most discordant scores between DMS methods in our meta-analysis that may reflect differences in the kinetics of RHO trafficking between the methods (see Discussion). Aside from T17M, which is an annotated dual Class 2 and Class 4 variant, no Class 1, 4, 5, 6, or 7 variants exhibited clear surface expression defects (**Figure 3A-B**), pointing to the specificity of both methods in measuring trafficking defects for Class 2 and, to a lesser extent, Class 3 mechanistic classes.

**Figure 3.**
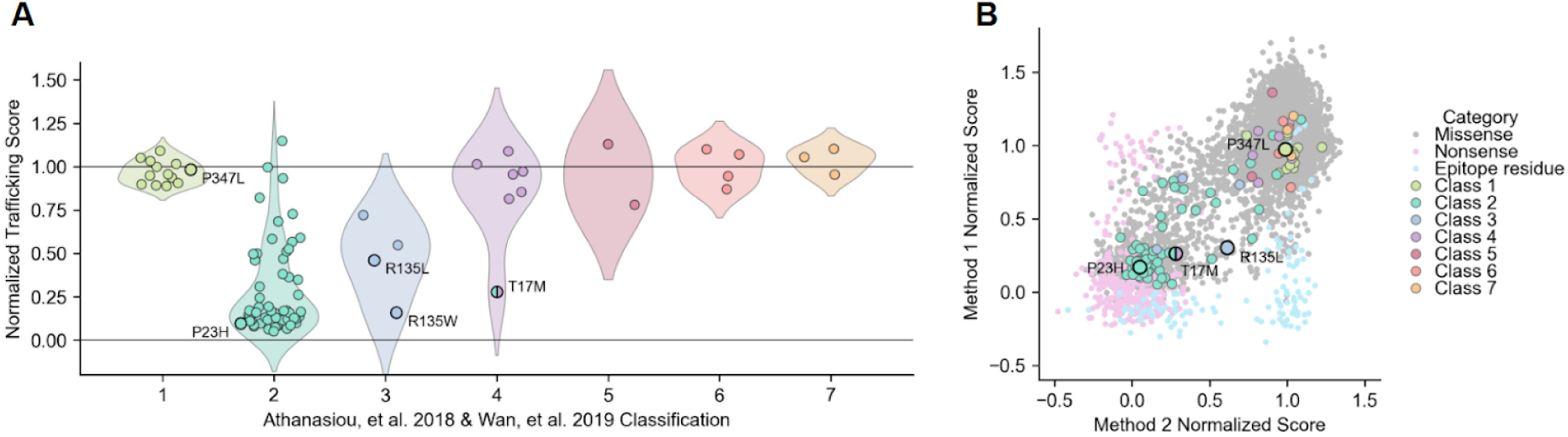
Variants with trafficking defects are enriched among the misfolding mechanistic class. (**A**) Distributions of the composite trafficking scores (y-axis) of literature-annotated variants stratified by their biochemical mechanistic class descriptions in Athanasiou, et al. 2018 and updated in Wan, et al. 2019. Composite trafficking scores are normalized from 0 (mean of non-trafficking negative controls) to 1 (wild-type surface trafficking, positive control). The biochemical mechanistic classes are as follows: (1) failure to localize to the outer segment, (2) misfolded and trapped in the ER, (3) defective endocytosis, (4) defective post-translational modification, (5) defective G protein activation, (6) constitutive activation, and (7) defective dimerization. Select variants are indicated. (**B**) Scatterplot showing trafficking scores from Method 1 (y-axis) and Method 2 (x-axis) at the variant level overlaid with variant classifications as reviewed in Athanasiou, et al. 2018 and updated in Wan, et al. 2019. Select variants are labeled: Class 1 - P347L, Class 2 - P23H, Class 3 - R135L, dual Class 2/4 - T17M.

### Evaluating trafficking assay sensitivity for detecting *RHO* variants

Because the trafficking assay was primarily designed to identify Class 2 (misfolding and mistrafficking) *RHO* variants, we compared our assay results with ClinVar annotations to see which known variants were identified as having pathogenic trafficking levels. ClinVar is a well-established database of genetic variants categorized as benign, pathogenic, or VUS based on the body (or lack thereof) of functional, genetic, clinical, and other types of evidence (https://www.ncbi.nlm.nih.gov/clinvar/). First, we implemented numerical cutoffs for ‘moderate confidence’ (< 0.7) and ‘high confidence’ (< 0.5) surface trafficking defects (using the upper limit of the 95% confidence interval) as well as ‘low’ (< 0.5) and ‘very low’ (< 0.25) surface trafficking (using the mean) based on the composite trafficking score of known benign variants and Class 2 variants (**Figure S3**). Then, we compared our assay results with ClinVar annotations. Approximately 55% of variants in the combined pathogenic and likely pathogenic categories exhibited moderate or high confidence surface trafficking defects, while no variants in the combined benign and likely benign categories did (**Figure 4A**). Thus, while a trafficking defect strongly suggests pathogenicity, a normal score does not exclude it, as six other proposed disease mechanisms exist^24^. In other words, the surface trafficking assays have excellent specificity and moderate sensitivity when predicting pathogenicity. In this context, the subset of VUS that exhibit moderate or high confidence trafficking defects now have data from a well-validated functional assay that supports their reassignment towards likely pathogenic or pathogenic categories (**Figure 4A**)^20^. Of the 6,612 missense variants, 757 (11.4%) had moderate or high confidence trafficking defects, substantially increasing the number of variants with evidence supporting pathogenicity compared to the 198 likely pathogenic or pathogenic variants currently in ClinVar.

**Figure 4.**
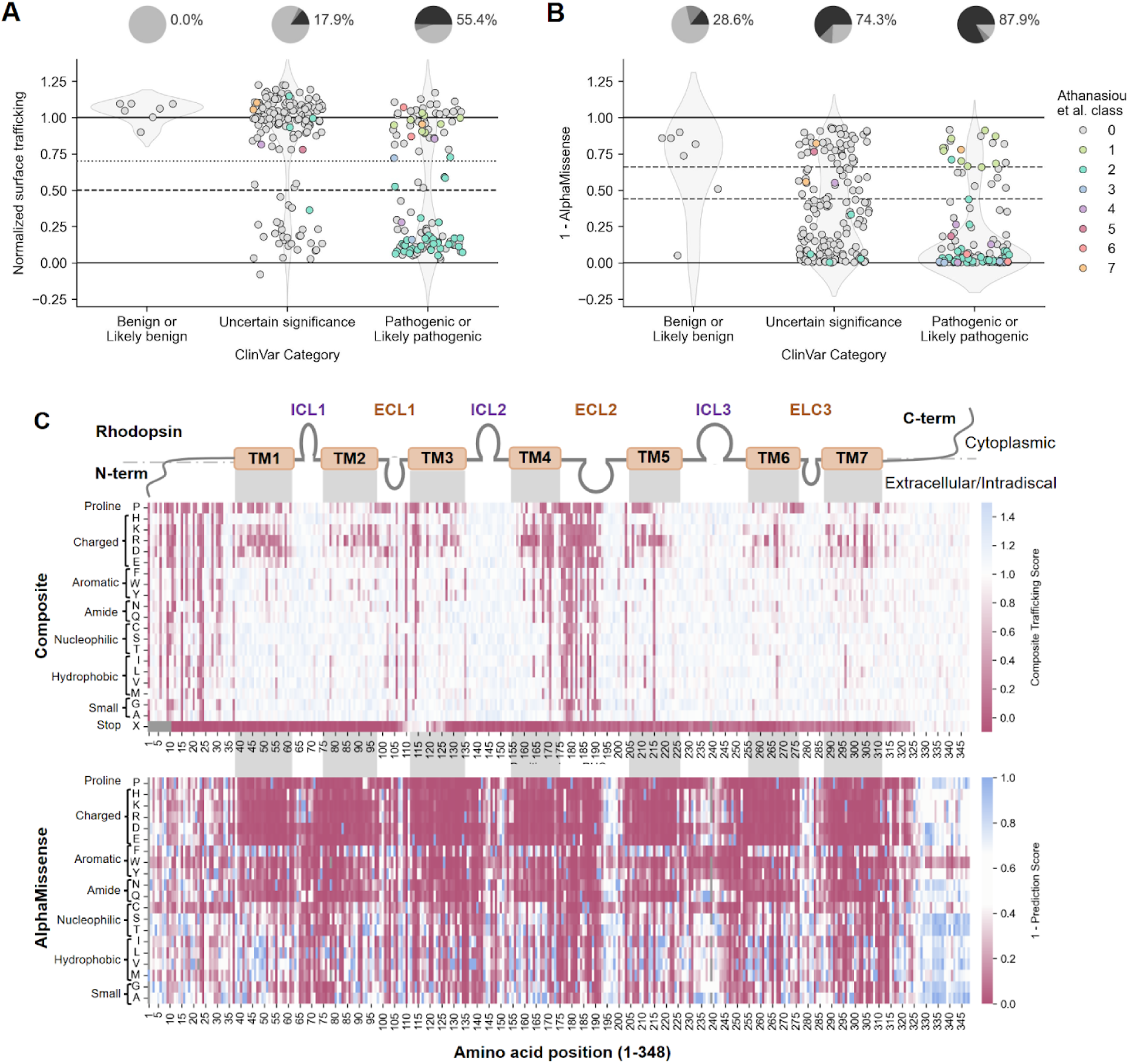
Trafficking defects predict variant pathogenicity. (**A**) Strip plot showing the composite normalized trafficking score for Method 1 and Method 2 for variants registered in the ClinVar database, stratified by clinical significance and colored by mechanistic classification as described in Athanasiou, et al. 2018 and Wan, et al. 2019 (0 = unclassified). The dotted horizontal line at y = 0.7 represents the threshold for moderate confidence trafficking defects, while the dashed horizontal line at y = 0.5 represents the threshold for high confidence trafficking defects. Pie charts placed above each category show the percentage of variants with moderate or high confidence trafficking defects (light gray = no defect, medium gray = moderate confidence defect, dark gray = high confidence defect). (**B**) Strip plot showing one minus the AlphaMissense pathogenicity prediction score (0 = strong prediction of pathogenicity, 0.44-0.66 = uncertain prediction, 1 = strong prediction of benignity), stratified by clinical significance and colored by mechanistic classification, for the same set of variants as in (A). (**C**) Heatmaps showing composite trafficking scores (top) versus one minus prediction score from AlphaMissense pathogenicity prediction score (bottom). The x-axis indicates the position along the RHO protein, with domain boundaries indicated (TM = transmembrane, ICL = intracellular loop, ECL = extracellular loop). The y-axis shows the amino acid substituted at each position.

### Empirical versus computational predictors of pathogenicity

Computational predictors, such as AlphaMissense, have been increasingly used to infer variant pathogenicity, guiding both genetic interpretation and early-stage drug discovery efforts. By leveraging evolutionary conservation and structural information, these models aim to classify missense variants as benign or pathogenic, often serving as a filter for identifying disease-associated alleles and for prioritizing therapeutic intervention targets. To assess how well AlphaMissense predicts pathogenicity for *RHO* variants, we compared the AlphaMissense predictions to the known ClinVar clinical significance category. Of the seven likely benign or benign *RHO* missense variants, AlphaMissense predicted one as ambiguous and another as likely pathogenic (**Figure 4B**), in contrast to our trafficking scores, which did not identify defects for any benign variants (**Figure 4A**). AlphaMissense called many more variants as pathogenic, including the majority of VUS and pathogenic *RHO* missense variants and many mistrafficking variants; however, it failed to predict pathogenicity for Class 1 variants, in the C-terminal tail. Compared to our composite trafficking score, AlphaMissense undercalls mistrafficking variants in the unstructured N-terminus and likely inaccurately predicts very high pathogenicity rates for the majority of uncharged transmembrane variants and tryptophan and cysteine substitutions (**Figure 4C**).

We evaluated 53 other computational metrics as well, demonstrating that the trafficking score provides distinct information not captured by the other known predictors (**Figure S4**). These results highlight that existing *in silico* tools can exhibit biases in pathogenicity predictions, often fail to offer the mechanistic detail necessary for guiding precision drug development, and would benefit from the integration of functional data into training datasets to improve accuracy.

### Broad correction of variant trafficking defects by a non-retinoid pharmacological chaperone

Protein misfolding diseases, where protein variants fail to fold and do not localize to the proper subcellular compartment, represent a significant therapeutic challenge. However, pharmacological chaperones, or ’correctors,’ offer a promising approach by stabilizing misfolded proteins and promoting their proper trafficking. Successful corrector therapies, such as ivacaftor for CFTR in cystic fibrosis^35^ and migalastat for GLA in Fabry disease^36^, have demonstrated the clinical potential of this strategy. *RHO*-adRP is particularly well-suited for a corrector therapy, as many pathogenic *RHO* variants lead to protein misfolding and retention in the ER. Previous work has explored retinoid analogs of the natural RHO ligand, 11-cis-retinal, which have shown low levels of corrector activity for variants like P23H^37–39^ but are light-sensitive and covalently conjugate to RHO. A recent study identified a non-retinoid pharmacological chaperone, YC-001, that reversibly binds the same orthosteric site for enhanced and more potent restoration of folding for four Class 2 variants with baseline trafficking defects, including P23H, D190N, G106R, and P267L^27^.

To investigate the corrective potential of YC-001 more broadly across mistrafficking variants, we performed the Method 2 DMS assay in the presence of 30 μM YC-001, the approximate EC_90_ concentration for P23H trafficking rescue (data not shown). Of the 1,260 moderate and high confidence mistrafficking variants identified by Method 2 at baseline conditions in the absence of corrector drug, 642 variants (51%) showed significant rescue (non-overlapping 95% confidence intervals) after 24 hours with 30 μM YC-001 (**Figure 5A,E**). This included most N-terminal and many transmembrane and ECL2 mistrafficking variants (**Figure 5B-D**). Particularly mutationally intolerant variants included those at C110 and C187, which form a disulfide bridge critical for folding and functioning of rhodopsin^29^, as well as other residues within ECL2 that may be important for coordinating protein-drug interactions (**Figure 5A,F**). Only one variant, A132C, which has no baseline trafficking defect, exhibited a significant (but small) decrease in trafficking with YC-001 treatment. Among the most common misfolding and mistrafficking variants identified in clinical cohorts in the United States, the majority of variants exhibited significant increases in trafficking with YC-001 treatment (**Figure 5F**), suggesting a substantial fraction of *RHO*-adRP cases caused by misfolding and mistrafficking variants may be addressable by small molecule correctors.

**Figure 5.**
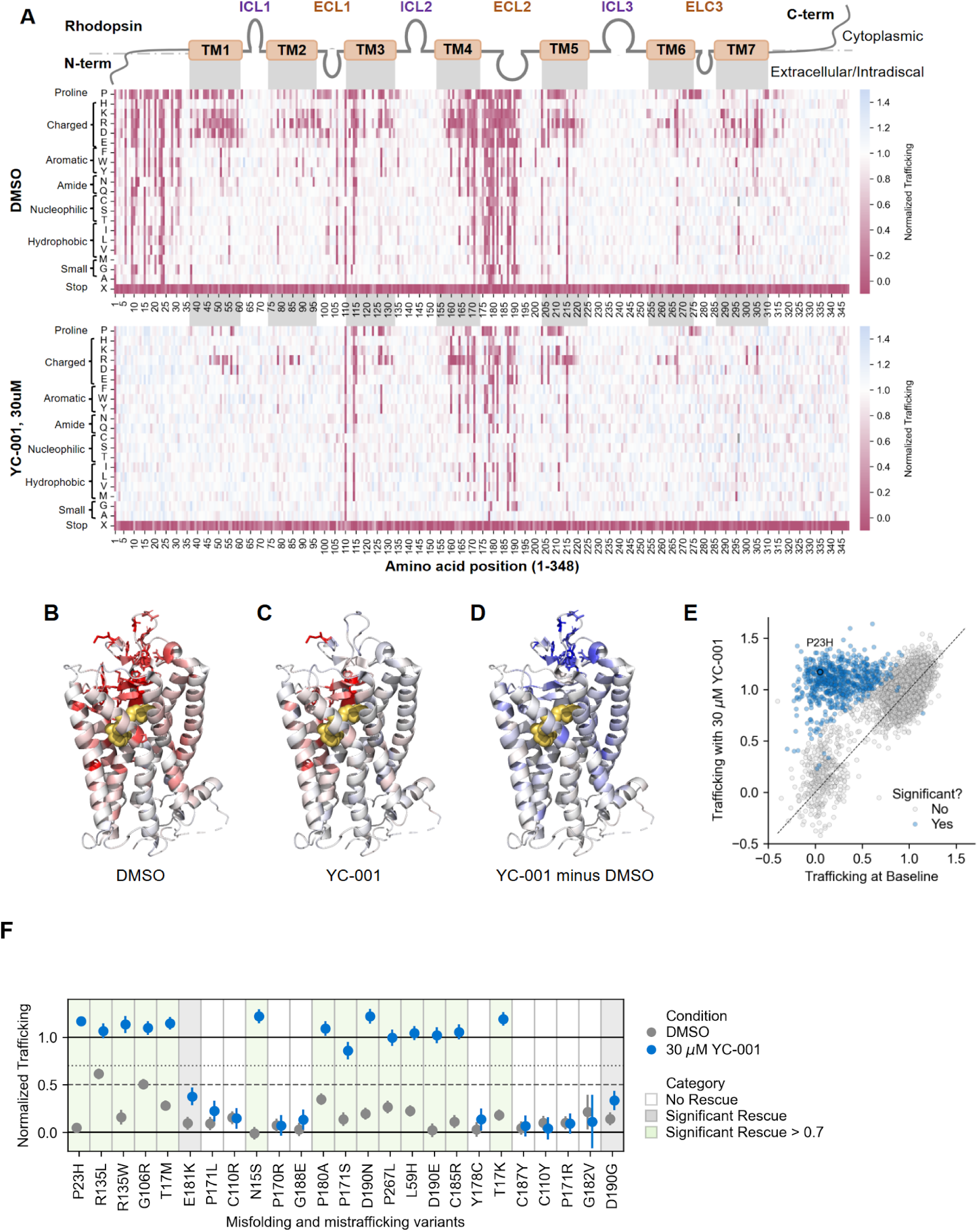
YC-001 broadly rescues trafficking for a majority of misfolding and mistrafficking variants. (**A**) Heatmaps showing surface trafficking for Method 2 at baseline with the DMSO vehicle (top, same as shown in Figure 2D bottom) and with 30 μM YC-001 treatment (bottom) for all missense and nonsense variants. The x-axis indicates the position along the RHO protein, with domain boundaries indicated (TM = transmembrane, ICL = intracellular loop, ECL = extracellular loop). The y-axis shows the amino acid substituted at each position. RHO surface trafficking is normalized on a log scale from 0 (mean of non-trafficking, nonsense variant negative controls) to 1 (wild-type RHO surface trafficking, positive control) and is indicated by cell color, with white representing trafficking similar to wild-type, pink representing a trafficking defect relative to wild-type, and blue representing a trafficking increase relative to wild-type. Missing measurements are indicated in gray. (**B-C**) RHO structure (PDB: 1F88) colored according to the average trafficking score at each position (excluding nonsense variant) with (B) DMSO vehicle or (C) 30 μM YC-001. More red represents a greater average trafficking defect. (**D**) RHO structure is colored according to the difference in average trafficking scores at each position between 30 μM YC-001 and DMSO vehicle. More blue represents a greater average trafficking rescue. (**E**) Scatterplot comparing trafficking scores at baseline with the DMSO vehicle (x-axis) versus with 30 μM YC-001 (y-axis). Variants with significant differences between treatment conditions (non-overlapping 95% confidence intervals) are shown in blue, with 643 variants above the y = x line (increased trafficking with 30 μM YC-001) and 1 variant below (decreased trafficking). (**F**) Rescue plots showing surface trafficking at baseline with DMSO vehicle (gray) and with 30 μM YC-001 (blue) for higher-frequency misfolding and mistrafficking variants identified in clinical cohorts, ordered by relative frequency. Trafficking is shown as the mean and 95% confidence interval. Thresholds for moderate confidence (< 0.7) and high confidence (< 0.5) trafficking defects are indicated as dotted and dashed lines, respectively. Variants that show no rescue have a white background. Variants that show significant rescue (non-overlapping confidence intervals) to a level below 0.7 have a gray background. Variants that show significant rescue to a level above 0.7 have a green background.

## Discussion

In this study, we used two complementary deep mutational scanning (DMS) approaches in HEK293T cells to systematically measure *RHO* variant effects on surface trafficking and identify those with pathogenic levels of misfolding and mistrafficking. Method 1 was a flow cytometry-based immunoassay similar to previous studies, while Method 2 was a proximity-based transcriptional reporter assay. The high correlation between the methods gives rigor to the interpretation of the results when low trafficking was observed in both assays, as indicated by the “high confidence” flag. Small areas of disagreement were expected from technical artifacts, such as within the antibody binding epitope (residues 2-10) in Method 1. These data agree well with prior studies conducted at lower throughput, especially those using flow cytometry assays similar to Method 1^24–26,40^. Overall, these data represent a major expansion of knowledge about sequence variation in *RHO*; while over 30 years of *RHO* research have identified 198 pathogenic or likely pathogenic human variants of any type (ClinVar, accessed February 2025), the data from this study identified over 700 missense variants with pathogenic trafficking levels. Moreover, 51% of these variants showed a significant increase in trafficking after treatment with YC-001, demonstrating the potential for small molecule corrector therapy for *RHO* variants. Altogether, these data demonstrate how DMS approaches can rapidly identify variants with functional defects and inform how to correct them, at scale.

Our assays reproducibly measured surface trafficking defects for most annotated and functionally validated Class 2 variants^10,24^, which induce misfolding and retention in the ER^41^. Empirically, low trafficking levels were detected for most misfolding variants (Class 2), some endocytosis variants (Class 3), and one post-translational modification variant (Class 4), but did not detect trafficking defects in Classes 1, 5, 6, 7 (outer segment trafficking, transducin binding, constitutive activation, and dimerization)^10,41^. Therefore, a normal (high) trafficking score does not guarantee that a variant is benign. Additional assays will be needed to detect variants in these classes. Until that time, when used as an assay to identify pathogenic variants, the trafficking score data in this study should be considered to have high specificity but imperfect sensitivity. A noticeable class with many undetected variants is Class 1, affecting the VxPx motif in the C-terminal tail^10^. Even so, we speculate that all missense variants in the other classes will be far fewer than the 757 mistrafficking variants identified here. It may be the case that *RHO*-adRP is primarily a misfolding disease^42^, but with some exceptions known due to its long history of investigations. This observation suggests the utility of focusing on “correctable” versus “uncorrectable” variant classes (or responder / non-responder) in the context of particular investigational therapies.

Regarding the “cutoff” value of the composite trafficking score which represents a pathogenic variant, it was notable that there was a bimodal distribution of trafficking scores (**Figure 2A,C,F**). Because of that distribution, different choices for the cutoff score for determining a pathogenic trafficking level affect the relatively small number of points in the middle of the distribution (**Figure S2**). For example, 757 of the 6612 variants (11.4%) showed trafficking scores statistically lower than a normalized composite trafficking score of 0.7, while 778/6612 (11.8%) variants had scores <0.5 using a hard cutoff. One helpful reference is that none of the 7 known benign or likely benign missense variants had a trafficking score below 0.899 (**Figure S3**). Nonetheless, future work could refine these cutoffs based on additional biological and genetic data, or implement a probabilistic / Bayesian framework.

The small number (14) of classically-described Class 2 variants that did not have low trafficking scores in this study (or in similar assays^24^) could be explained by a number of reasons, including: differences in the rhodopsin sequence used in earlier research (e.g. bovine backbone)^43^; differences in protein degradation or posttranslational modification in different cell types^44^; differences in folding, maturation, and trafficking through the secretory pathway^45^ (though these are thought to be universal^46^); or they truly were not mistrafficking variants.

Similarly, further examination of the rare instances of disagreement between Method 1 and Method 2 may provide additional insight into variant behavior. For example, the class 3 variants, affecting the conserved arginine 135 in the E/DRY motif^47^ and associated with altered endocytosis^48^, exhibited a range of trafficking defects. The most severely affected, R135W, has been described as having Class 2 characteristics^47,49,50^, unlike the other Class 3 variants R135G, R135L, and R135P^51^. Differences in the kinetics of Method 1 and Method 2 could explain the lower trafficking scores of Class 3 variants in Method 1. Method 1 measured steady-state RHO surface abundance, while the proximity sensor in Method 2 may be less affected by surface residence time. R135L is normally hyperphosphorylated^51^, bound to arrestin, and endocytosed, which could explain the observed lower score by Method 1 and higher score by Method 2. At this amino acid position and others, DNA sequencing errors and coverage issues could be a source of assay disagreement, though we speculate this would be very rare.

Regarding comparing empirical DMS data to computational predictors, it was notable that even a modern algorithm like AlphaMissense^52^ did not perform as well as the trafficking score in predicting pathogenicity (**Figure 4**). This supports the premise that empirical cell-based assays add a great deal of information beyond what computational predictions can currently provide. In fact, empirical data from DMS experiments could be a valuable addition for re-training the machine learning based computational models to improve their accuracy.

Decades of research on *RHO* variants have identified the structural constraints of RHO with its bound cofactor, 11-cis retinal^53,54^. With the comprehensive overview provided by this data highlighting regions where variation causes misfolding and mistrafficking (**Figure 2D-F**), two major patterns emerge. First, the region of highest constraint is located above the retinal binding pocket in the extracellular/intradiscal space, confirming and extending earlier data^55^ that the “beta-plug” region is critical for RHO protein folding and stability. The second broad pattern that emerges is that transmembrane domains are susceptible to charged and proline amino acid substitutions, as expected^56–58^. Other notable constrained residues are cysteines involved in disulfide bridges. Compared to a DMS dataset for the beta-adrenergic receptor^56^, another class A GPCR, the datasets have localized constrained residues in common, such as the (Wxx)GxxxC “structural hinge” motif at the beginning of TM3. However the striking, broad mutational intolerance in the N-terminal and ECL2 regions, around and forming the beta-plug, are not found in the beta-adrenergic receptor, highlighting the distinctive mutational sensitivity of the extracellular/intradiscal region of RHO. One explanatory hypothesis is that opsins do not bind their ligands through “open” extracellular domains present in most GPCRs but instead bind the retinal ligand through transient pores that open between transmembrane helices, allowing the extracellular/intradiscal region to evolve to contribute to thermal stability and receptor activation^10,59^. Consequently, compared to other GPCRs, the unique way opsins bind to their ligand may create opportunities for tailored drug design.

*In vitro* studies to date have shown proof-of-concept for multi-variant, non-retinoid pharmacological chaperones of RHO, with a recent study demonstrating variable trafficking rescue for a small subset of 123 pathogenic missense variants^60^. Using our high-throughput approach, we identified broad variant coverage (51%) for the well-established research corrector compound, YC-001^27^, with the RHO N-terminus and ECL2 being generally both highly mutationally sensitive and highly amenable to treatment. The rarer, non-responsive variants included C110 and C187, which are involved in a structurally important disulfide bond^61^. While the micromolar potency of YC-001 and its partially-characterized pharmacokinetic properties preclude its direct development into a therapy, our data underscore the potential of an orally bioavailable, broad-spectrum, non-retinoid corrector. Such a treatment could address the majority of *RHO*-adRP cases driven by protein misfolding while retaining the advantages of oral delivery. Furthermore, this study presents a scalable and repeatable strategy for assessing the variant landscape at baseline and in the presence of small molecule corrector candidates for protein misfolding diseases more broadly.

In summary, multiplexing of a biologically straightforward protein trafficking assay across a collection of every missense variant efficiently produced a large volume of structural and pharmacogenomic information about *RHO*, an important inherited retinal disease gene. This study greatly expanded the knowledge of *RHO* variation, more than tripling the number of variants with pathogenic trafficking levels compared to prior knowledge of pathogenic human variants from decades of prior research. Each of the mistrafficked variants can be classified as responding or not responding to a small molecule corrector, and most mistrafficking variants did respond to corrector therapy. Experiments are ongoing both to develop next generation corrector molecules as well as developing assays to increase the sensitivity of RHO DMS assays. This approach also demonstrates how multiplexed methods can be used to efficiently implement a comprehensive approach for both diagnosing and treating genetic diseases.

## Supplementary Figures

**Figure S1.**
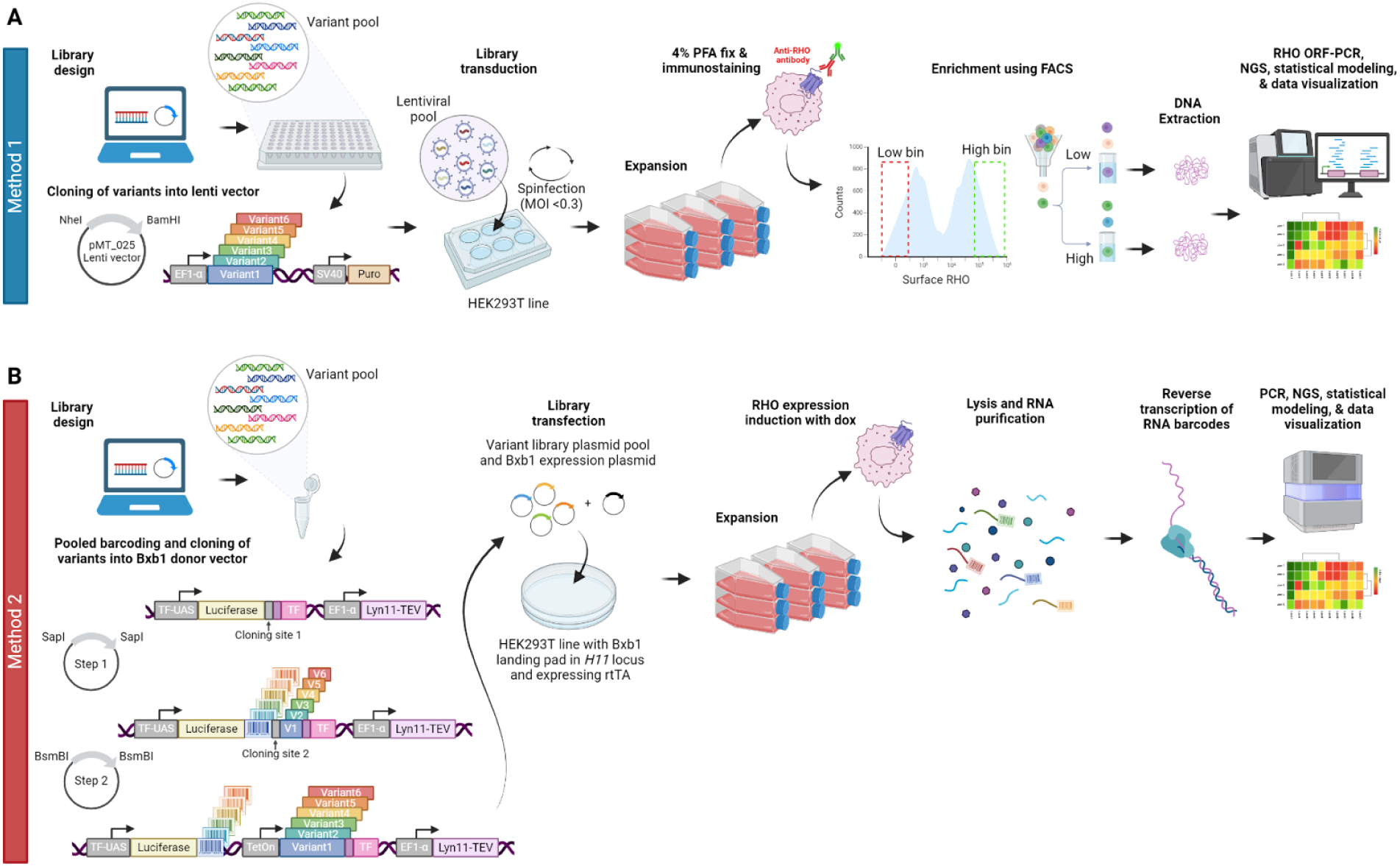
Detailed schematic of deep mutational scanning workflows. **(A)** Method 1 involves *in silico* library design, cloning variants into the pMT_025 lentiviral vector (NheI/BamHI = restriction sites, EF1α/SV40 = strong promoters, Puro = puromycin resistance gene), generating a lentiviral pool (MOI = multiplicity of infection), and transducing HEK293T cells at single copy. For the assay, cells are expanded, fixed with paraformaldehyde (PFA), and immunostained for surface-displayed RHO. The top and bottom quartiles are isolated by fluorescence-activated cell sorting (FACS) for downstream DNA extraction and library preparation (ORF-PCR = open-reading frame polymerase chain reaction). Libraries are subjected to next-generation sequencing (NGS), statistical modeling is performed on the sequencing counts to infer variant effects, and data are further analyzed and visualized. **(B)** Method 2 involves *in silico* library design, a multi-step variant barcoding and cloning process into a Bxb1 integrase-compatible donor vector, and transfecting this plasmid library along with a Bxb1 integrase expression plasmid into a HEK293T cell line that (i) harbors a landing pad for single-copy, site-specific integration and (ii) expresses the reverse tetracycline transactivator (rtTA) for doxycycline (dox)-inducible activation of the TetOn promoter that drives expression of RHO variants fused to a transcription factor (TF) via a TEV protease-cleavable linker. For the assay, cells are expanded and RHO expression is induced with dox. A TEV protease anchored to the plasma membrane via the Lyn11 domain cleaves the linker between properly trafficked RHO and the TF, allowing for the TF to translocate to the nucleus, bind a TF upstream activating sequence (UAS), and induce expression of a reporter gene with a *RHO* variant-specific RNA barcode. Cells are lysed, RNA is purified, and barcode RNA is selectively reverse transcribed and amplified with PCR. Libraries are subjected to NGS, statistical modeling is performed on the sequencing counts to infer variant effects, and data are further analyzed and visualized.

**Figure S2.**
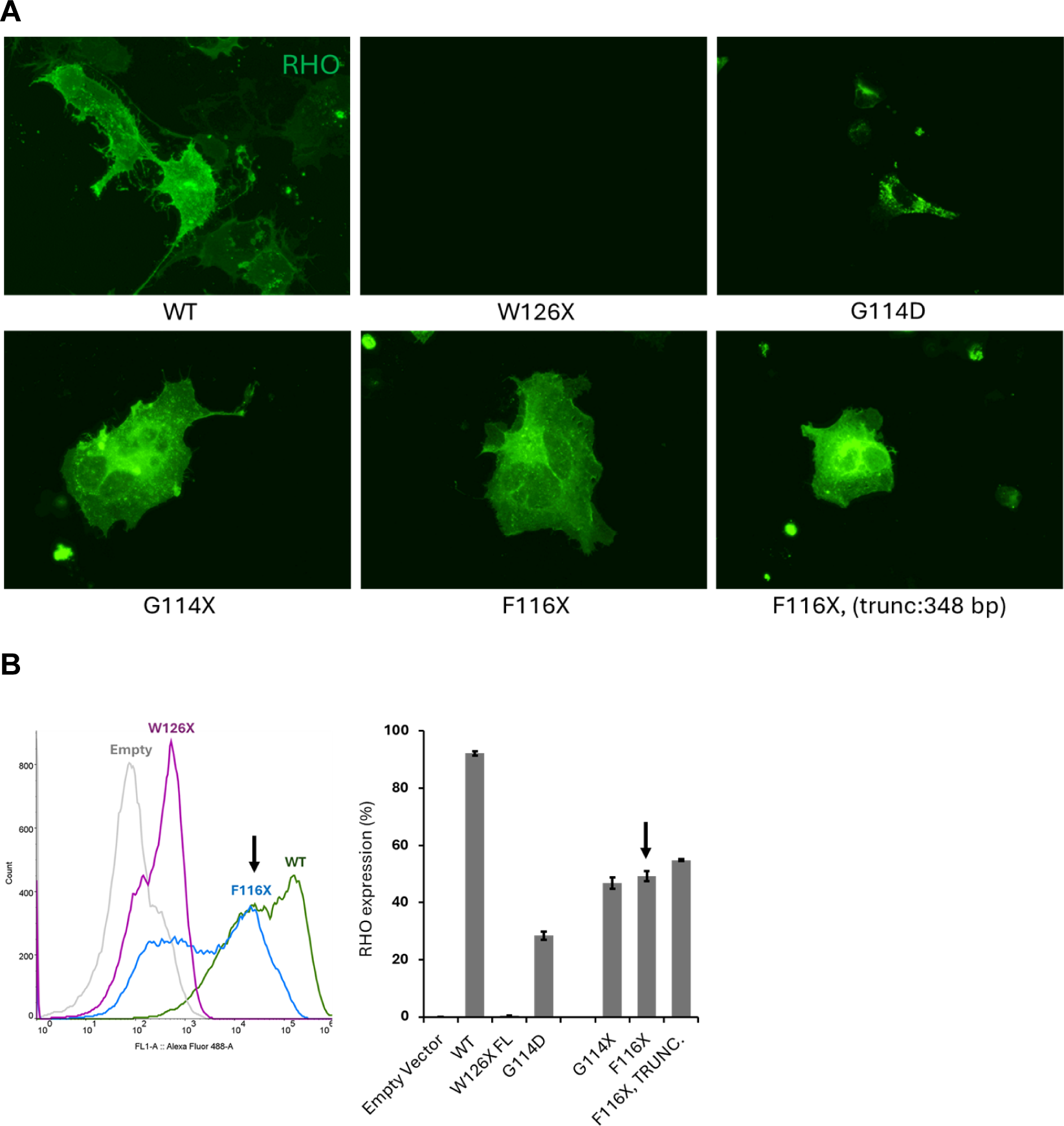
Nonsense variants between residues 110-125 show surface localization in transfected HEK293T cells. In contrast to WT rhodopsin, nonsense variants in the DMS library categorized as ’X’ (in the heatmap) were predicted to have low surface trafficking scores. However, nonsense variants in the amino acid sequence between positions 110-125 exhibit intermediate-to-high surface trafficking scores in Method 1. To further investigate these findings, two nonsense variants from this region, and appropriate controls, were transfected and characterized by immunofluorescence and flow cytometry. In this experiment, nonsense variants G114X, F116X, and W126X (designed to have full-length cDNA sequences) and a nonsense variant F116X trunc. (with a shorter cDNA, with sequence termination after the stop codon) were used. **A)** Fluorescence microscopy images shows surface localization of RHO protein, with G114X, F116X, F116X truncated variants, alongside wild-type, in transfected HEK293T cells. In contrast, a nonsense variant W126X fails to form functional protein due to a PTC (premature termination codon), and a known class II misfolding variant G114D shows low surface expression. **B)** Flow cytometry analysis indicating nonsense variants G114X, F116X, and F116X (trunc.) with intermediate RHO cell surface trafficking. Wild-type RHO served as a positive control, exhibiting robust trafficking. In contrast, W126X displayed no detectable surface expression, while G114D showed reduced surface expression (n=3). Both assays were performed on non-permeabilized cells. (Empty vector = backbone vector with no insert, WT = Wild-type rhodopsin, G114X, F116X, and F116X (trunc.) = a nonsense variant with surface trafficking, W126X = a nonsense variant with no surface trafficking, G114D = a class II variant.

**Figure S3.**
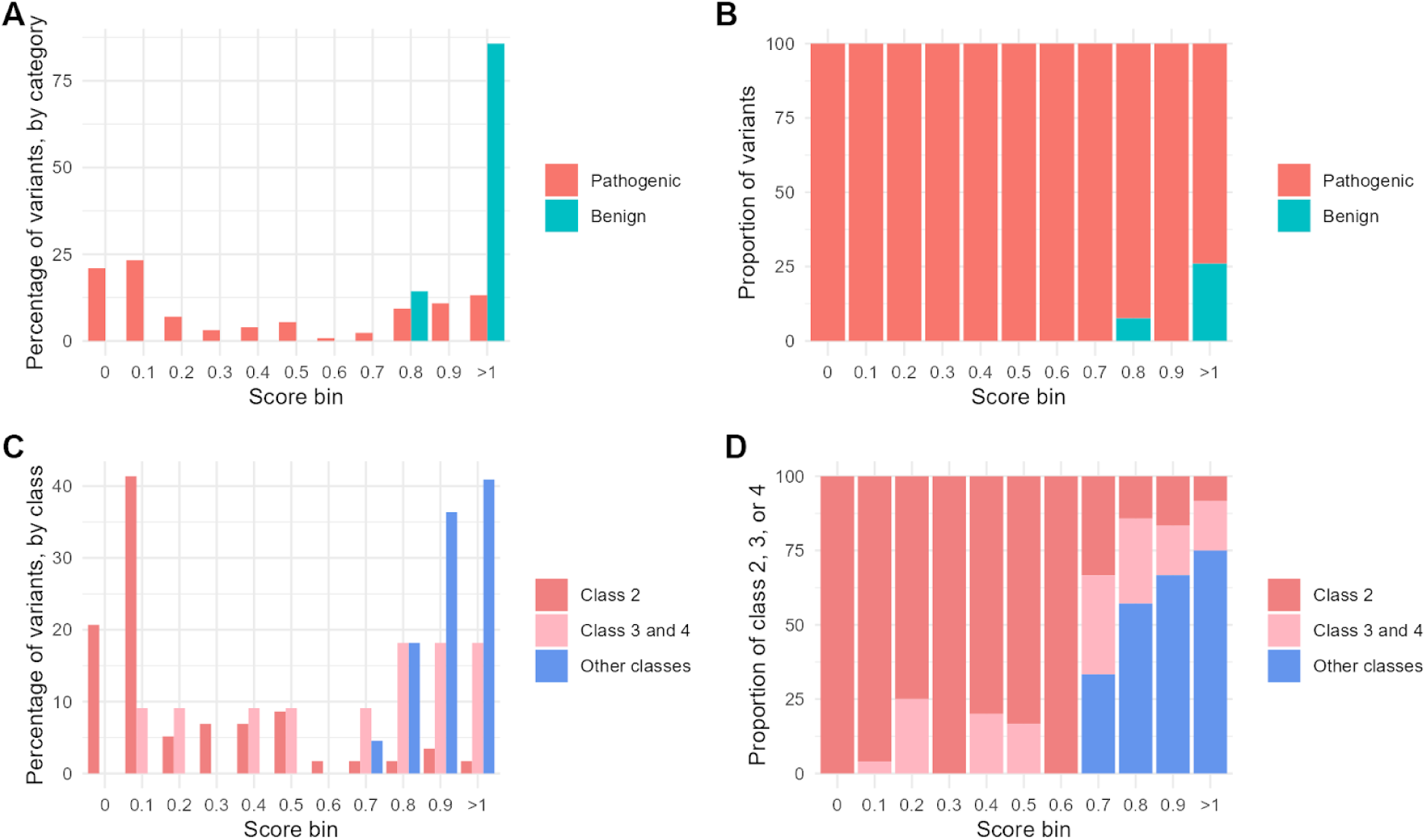
Investigating pathogenicity cutoff values. To determine which trafficking scores correspond to pathogenic variants, the ClinVar pathogenicity (top) or rhodopsin mechanistic classes (bottom) are shown as a function of trafficking score bins. Results are shown with two different normalization approaches: as a percentage of variants in each category (left, **A, C**), or as a proportion of between the two categories (right, **B, D**). There are no benign variants or non-class 2,3,4 variants with a trafficking score < 0.7, which defined the cutoff for moderate confidence mistrafficking variants. A cutoff for high confidence mistrafficking variants was set at 0.5 because the underlying data is sparse (top: 7 of 136 variants are benign; bottom: 22 of 144 variants belong to the Other classes).

**Figure S4.**
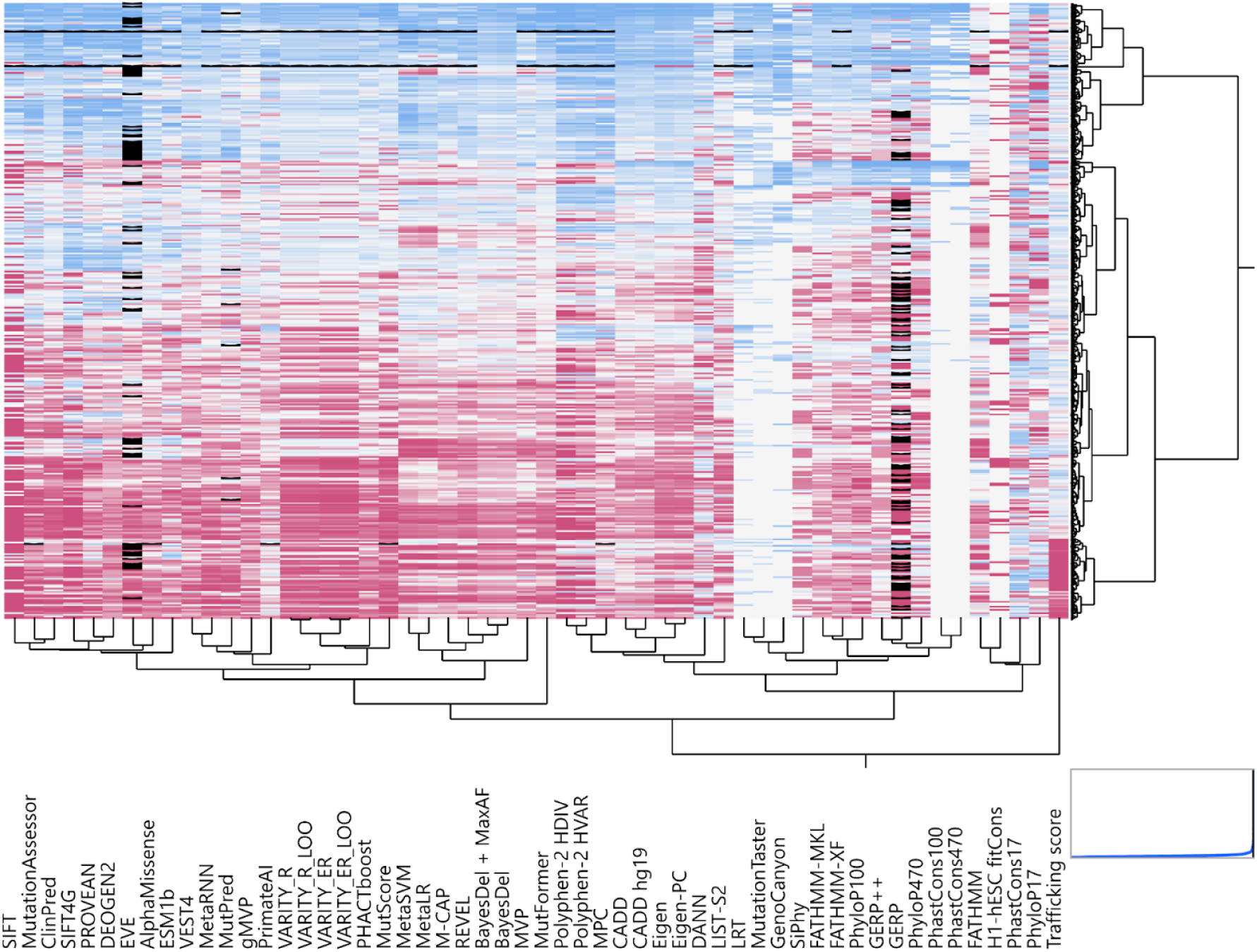
Comparison of trafficking scores with computational predictors of pathogenicity. (**A**) Hierarchical clustering groups variants (rows) and predictors (columns) that display similar patterns. Blue (0) represents a benign prediction while Red (1) represents a pathogenic prediction. The trafficking score has the longest dendrogram branch, indicating it is the most distinctive predictor. Hierarchical clustering was performed using the Ward method with robustly normalized columns, missing value imputation (black), and with row clusters ordered by the first principal component of the data (JMP software v17.2).

## Methods

### Cell culture for Method 1

For Method 1, HEK 293T cells (ATCC, CRL-3216) were maintained in Dulbecco’s modified Eagle’s medium (DMEM) (Thermo Fisher, 11-995-065) supplemented with 10% fetal bovine serum (Thermo Fisher, Cat.SH3007103). The cells were maintained at 37°C in 5% CO_2_.

### Vectors and library preparation for Method 1

For the design of the saturation mutagenesis variant library, each amino acid position (a.a. 2-348) in the reference sequence of *RHO* was systematically altered to 19 different amino acids, as well as synonymous and stop codons that serve as assay controls. The wildtype human *RHO* sequence was based on the open reading frame (ORF) in NM_000539.3. Variant codons (usually 1 per amino acid position) were selected to favor 2- or 3-nucleotide changes rather than 1-nucleotide changes when possible, to avoid collisions with sequencing errors. Variant codons which would create restriction sites used in the cloning reaction were excluded as well.

For synthesis of this library (Site Saturation Variant library product; Twist Biosciences), an error-free wild-type Rhodopsin template was synthesized as the starting point for all variants in the library. This WT template DNA was divided into aliquots and placed in individual wells of 96-well plates. To each well, outer primers were added, flanking the specific region of the gene where the variant would occur, enabling site-directed mutagenesis. PCRs were performed using high-fidelity polymerase to ensure the variant was incorporated into the newly synthesized DNA according to manufacturer’s instructions (Twist Bioscience). After PCR, the DNA from each well was normalized to ensure equal representation of all variants in the final library. The DNA from all wells was pooled together, and quality control (QC) was performed by NGS to verify the presence and accuracy of the desired variants. The ORF was flanked with adapters suitable for restriction digestion and ligation into NheI and BamHI sites of pMT_025 vector (Addgene #158579). The pMT_025 was provided by the Genetic Perturbation Platform, Broad Institute of MIT and Harvard and contains two components: 1) an EF-1α promoter that drives the rhodopsin ORF and 2) an SV40 promoter that drives puromycin resistance. The fragments were ligated into the NheI and BamHI restriction site of pMT_025 vector using T7 DNA ligase. Post clean-up, the products were transformed, and the plasmid DNA library was isolated using the Maxi-prep kit (Qiagen HiSpeed Maxi, 12663) and subjected to NGS. The mutagenesis library was characterized to identify and distinguish intended variants from variants arising from errors during library synthesis. To assess clone distribution, we scaled raw variant counts in the plasmid library by their local read coverage, and the raw counts were adjusted and scaled to account for uneven sequencing depth. The scaled values were log-transformed for better visualization and the resulting values were log_2_-transformed to form the histogram bins. Briefly the steps involved trimming the flanking sequences, aligning overlapping paired-end reads, applying quality and length filters, and identifying and quantifying sequence variation and the abundance of each variant. The variant abundance distribution revealed a representation of >99% of the intended substitutions in the variant pool.

### Lentivirus production for Method 1

HEK293T packaging cells were cultured in DMEM (Thermo fisher, 11965092) supplemented with 10% FBS (Hyclone, SH30071.03). Briefly, the cells were transfected with the plasmid DNA lentiviral library, psPAX2 (Addgene #12260), and pMD2.G (Addgene #12259) using TransIT-LT1 transfection reagent (Mirus Bio, MIR 2304). Details of the protocol can be accessed at http://www.broadinstitute.org/rnai/public/resources/protocols. Media was replaced and viral supernatant was harvested at 24 and 48 hours. Serial dilutions of a standard and library were used to determine the viral titer in A549 cells. The transduced cells were selected using puromycin (1.5 µg/mL, Gibco, A1113803) for two days. The AlamarBlue assay was performed, and the standard curve method was used to determine the linear range, sensitivity of the assay, and the viral titer of the *RHO* library.

For library transduction, HEK293T cells were cultured and expanded in T225 flasks. Upon reaching 80% confluency (∼45 million cells), the cells were washed with PBS and dissociated using 0.25% trypsin. To ensure adequate coverage/representation of the *RHO* library, each variant was transduced into a minimum of 500 cells. Additionally, the cells were transduced at a low multiplicity of infection (MOI=0.3) to ensure single-copy transgene integration. This ensures the required coverage/representation of the library containing up to 7,299 variants, with an estimate of 500 transduced cells per variant. With a target cell transduction efficiency of <30%, a total of 14.7 million cells were transduced by combining 48.9 mL of viral supernatant (titer of 2.42 x 10^5^) and polybrene (8 µg/mL, Sigma, H9268) and plated 2 mL per well into a total of four 6-well plates. Subsequently, the plates were sealed and centrifuged at 2000g for 50 minutes and maintained at 37°C with 5% CO_2_ overnight. The following day the media was replaced with fresh media. After 48 hours, the cells were dissociated using 0.25% trypsin, pooled and re-seeded at a density of 5 x 10^6^ cells per T225 flask in 50 mL growth media (a total of ten flasks). Selection pressure was applied by adding Puromycin at a final concentration of 800 ng/mL. On day 5-6 transduction efficiency was estimated in this fraction by immunostaining followed by MACS quant. To determine the infection efficiency, 3.5 x 10^5^ transduced and non-transduced cells were plated with and without Puromycin (800 ng/mL). The transduction efficiency was estimated as the ratio of average cell count of puromycin^+^ cells/average cell count of puromycin^-^ cells.

### Immunostaining and FACS for Method 1

Cells were washed with 10 mL DPBS per T225 flask and dissociated using 0.25% trypsin. The flasks were gently tapped and as soon as the cells got detached, 10 ml complete medium was added to quench the reaction. The cells from all flasks were pooled, centrifuged at 1000 rpm for 5 minutes at room temperature. The cell pellet was washed with 10 mL of DPBS and centrifuged at 1000 rpm for 5 minutes. The pellet was resuspended and fixed in 10 mL (10 million cells/mL) 4% paraformaldehyde (Electron Microscopy Sciences, 15710) for 20 minutes at room temperature. After PBS wash, blocking was performed with 3% BSA/PBS for 10 minutes followed by incubation in RetP1 *RHO* primary antibody (Sigma, O4886, 1:2000 dilution) for 30 minutes at room temperature. The cells were collected by centrifugation at 1200 rpm for 5 minutes, washed with 10 mL PBS, followed by incubation with anti-mouse 488 secondary antibody (Thermo, A21121) for 30 minutes in the dark at room temperature. After incubation, the cells were washed with PBS and resuspended at a density of 1 x 10^7^ cells/mL in PBS. All the cells were passed through a 40 µm cell strainer (Corning, 352340) before acquisition and sorting using Sony Cell sorter machine (SH800 or MA900). The sorting utilized a 100-micron nozzle, with the sort mode set to purity. The flow-rate was maintained between 2500-3000 events/sec, and sample pressure was kept constant at 7 psi, and was held at 4°C. After defining the gates using positive and negative controls, cells were sorted from RHO^high^ and RHO^low^ expressing bins. The top 20-30% percentile of each bin, estimated to represent the entire variants, was sorted and collected separately. Using the MA900 sorter, 8.5 x 10^6^ cells RHO^low^ cells and 12.3 x 10^6^cells RHO^high^ cells were sorted. Using SH800 sorter, 4.5 x 10^6^ cells RHO^low^ cells and 6.6 x 10^6^ cells RHO^high^ cells were sorted. The sorted cells were pelleted down and subjected to DNA extraction.

### Extraction of DNA from FACS sorted cells for Method 1

The recommended reagent volume described below is for the extraction of genomic DNA from 1x10^6^ fixed cells. The volume was scaled up based on the number of cells sorted. For Method 1, the cells were lysed overnight in 200 µL Quick-extract buffer (Lucigen, NC0302740) supplemented with 3.75 µL proteinase K (Thermo, EO0491) at 60°C, 600 rpm. Post lysis, 50 µL of 5M NaCl (Thermo Fisher, AM9760G) was added and mixed thoroughly by inverting the tube 50 times and incubated on ice for 10 minutes. Following the incubation, 3.5 µL of RNAase A (Thermo, EN0531) was added and mixed thoroughly by inverting the tube 50 times and incubated for 10 minutes at room temperature. The samples were centrifuged at 12,000 rpm for 10 minutes at room temperature. The supernatant was carefully transferred into a fresh tube and mixed with equal volume of Isopropanol and mixed thoroughly by inverting 20 times and incubated for 10 minutes at room temperature. The tubes were left undisturbed, which allowed the phases to separate. The top phase was carefully aspirated and discarded. Equal volume of monarch DNA binding buffer (Monarch Kit, T1030S) was added and the samples were mixed thoroughly by inverting the tube 10-15 times and transferred into the Monarch columns. The columns were centrifuged at room temperature for 1 minute, discarding the flowthrough. The columns were washed with 500 μL DNA Wash Buffer and centrifuged at max speed for 1 minute. The columns were then transferred into a clean 1.5 mL microfuge tube. Elution buffer was added (30 µL) to the center of the matrix, incubated for 1 minute, centrifuged at max speed for 1 minute to elute DNA. The concentration was measured using Nanodrop and integrity was confirmed by electrophoresis by running 1 µg DNA on 1% TAE agarose gel.

### Assay deconvolution for Method 1

The rhodopsin ORF from the genomic DNA was PCR amplified using primers: Forward: 5′ ATTCTCCTTGGAATTTGCCCTT−3′, Reverse: 5′-CATAGCGTAAAAGGAGCAACA −3′. PCRprimers were designed to have ∼100 bp flanking sequences on either side of the ORF region and was performed using Q5 Hot Start High-Fidelity 2X Master Mix (NEB, M0494S) using program: 95°C for 30 seconds, 98°C for 10 seconds, 69°C for 30 seconds, 72°C for 2.5 minutes (35 cycles) and 72°C 2 minutes and 4°C hold. A full 96-well PCR plate was used for each set, and each well with a reaction mix of 50 µL and 250 ng of gDNA. Post-PCR, the products were separated on 1% agarose, extracted and purified first using QIAquick Gel Extraction Kit (Qiagen, 28704) followed by AMPure XP kit (Beckman Coulter, A63881). PCR products from each set were subjected to transposon-mediated fragmentation, indexed and sequenced on an Illumina next-generation sequencing platform. Screen deconvolution was performed with Analyze Saturation Mutagenesis (ASM) software, as described^62^. Briefly, paired-end reads were aligned to the reference sequence, filtered, trimmed, and variant counts were calculated and the output files were parsed, annotated, and counts were merged into a single .csv file.

### Statistical Analysis for Method 1

For both datasets, variant counts were modeled using the *glmmTMB* package with a negative binomial linking function, using R version (4.4.1). For Method 1, a mixed-effects model was fitted to each position to estimate the effect sizes of amino acid type, gate conditions, and amino acid and gate interactions, while accounting for replication effects. Estimated marginal means were calculated, and linear contrasts were performed as the final estimate of relative abundance, which is the difference between abundance in high and low gates. The results were min-max scaled to read depth, with the mean linear contrast estimates of wild type as the maximum and the mean linear contrast of nonsense variants as the minimum and this resulted in most relative abundance values falling between 0 and 1.

### Variant library design and oligonucleotide pool synthesis for Method 2

The UniProt protein sequence for the 348 amino acid (aa)-long human rhodopsin protein (P08100) was used as the reference for library design and codon optimized at the DNA level for ease of DNA synthesis and downstream molecular cloning. All single-residue missense and nonsense variants (6,960 in total at the protein level) were designed *in silico* using up to three of the most common codons where possible (https://github.com/octantbio/rho-dms). Only nucleotides within the codon of interest were changed except in cases where the substitution introduced a homopolymeric sequence that could interfere with DNA synthesis or a restriction site that could inhibit proper molecular cloning. The 348 amino acid (aa)-long sequence was divided into five chunks between 68 to 70 aa in length to be within current pooled oligonucleotide synthesis length limits (≤ 300nt) after the addition of sequences required for amplification and cloning. Oligonucleotides for all five chunks were ordered in a single oligonucleotide pool from Twist Bioscience, with each chunk containing unique flanking sequences. Upon arrival, the lyophilized oligonucleotide pool was resuspended in IDTE pH 8.0 (IDT, 11-05-01-09) to 10 ng/μL, aliquoted, and stored for long term storage at -80°C.

### Step 1 library cloning for Method 2

Each chunk was amplified in two rounds of PCR using NEBNext® Ultra™ II Q5® Master Mix (NEB, M0544X), 0.2 ng/μL oligonucleotide pool template, and 5 μM forward and reverse chunk-specific primers. The first PCR selectively amplified the chunk of interest from the oligonucleotide pool, while the second PCR appended a variable 21-nucleotide barcode sequence and constant region for downstream cloning steps to each chunk sublibrary (cycle numbers for amplification determined empirically by qPCR).

Chunk-specific base cloning vectors were designed to contain a pair of SapI restriction upstream of a constant portion of the *RHO* gene. Golden Gate cloning with SapI (NEB, R0569L), high concentration T4 DNA Ligase (NEB, M0202M), 0.5nM of pre-digested vector, and 0.5nM amplified insert was performed in T4 DNA Ligase Reaction Buffer (NEB, B0202S) to insert the barcoded variable regions into the appropriate base cloning vector using the following cycling conditions: 37°C x 5 min initial digestion, 65 cycles of 37°C x 5 min digestion followed by 16°C x 5 min ligation, 37°C x 10 min final digestion, 70°C x 20 min denaturation, and 10°C hold. Golden Gate reaction products were concentrated using the DNA Clean & Concentrator-5 kit (Zymo Research, D4004) and eluting in 6 μL fresh MilliQ water. Two microliters of this concentrated Golden Gate product were electroporated into 50 μL of Endura™ Electrocompetent Cells (Lucigen, 60242-2) in Gene Pulser/MicroPulser Electroporation Cuvettes with a 0.1 cm gap width (Bio-Rad, 1652089) using a Bio-Rad MicroPulser Electroporator (Bio-Rad, 1652100) on program “Ec.1” with voltage 1.8kV and time constants between 5.5-5.7 ms. Immediately following electroporation, 2mL of pre-warmed Recovery Media (Lucigen, 80026-1) were added to the cuvette to collect the electroporated bacteria, which were transferred to a 14mL round-bottom tube and shaken at 200 rpm at 37°C x 1 hr to allow recovery and expression of the plasmid-encoded kanamycin resistance gene. A portion of the recovery culture was serially diluted and plated on LB agar plates with 50 µg/mL kanamycin sulfate (Teknova, L1025) for estimation of electroporation efficiency and library diversity. The remaining recovery culture was inoculated into 50mL 2XYT Broth (Teknova, Y2140) with kanamycin and grown for 16 hours at 30°C. For electroporation reactions yielding greater than 1e6 colony forming units, plasmid libraries were purified from the liquid cultures using the ZymoPURE™ II Plasmid Purification Midiprep Kit (Zymo Research, D4201) according to the manufacturer’s instructions.

### Barcode-variant mapping for Method 2

Following Step 1 library cloning, each plasmid contained a designed variable chunk sequence adjacent to an unknown random barcode and separated by a pair of Esp3I restriction sites. To map the relationships between the barcodes and variable chunk sequences, this region on the plasmid libraries was amplified by PCR using the NEBNext® Ultra™ II Q5® Master Mix and 5 μM forward and reverse primers containing sequences for grafting to an Illumina flow cell (single indexing). Two technical replicate amplicon sequencing libraries were prepared for each chunk and were subjected to 2x150 paired-end sequencing using the 300-cycle NextSeq 2000 P3 Reagent kit (Illumina, 20040561) on an Illumina NextSeq 2000 instrument. Resulting raw BCL files were demultiplexed with bcl2fastq and processed as described in^63^. Briefly, barcode and oligo sequences were extracted, mapped against a reference of all designed *RHO* sequences, counted, and filtered. Each oligo-barcode pair was required to: (i) be the appropriate length, (ii) have >=3 supporting read counts, (iii) have a purity >= 0.75 (defined as the fraction of all reads from the barcode which support the pair). Each library chunk was mapped separately, and then all resulting chunk maps were concatenated to generate the final barcode oligo map. Across the 12 assay samples, the median number of quantified barcodes per variant was 57.5.

### Step 2 library cloning for Method 2

A second Golden Gate cloning step was performed to insert the remaining elements (regulatory elements and upstream constant portion of the rhodopsin sequence) into the Esp3I restriction sites between the barcode and variable chunk sequence. The elements to insert for each chunk were encoded in chunk-specific donor vectors, and both these donor vectors and the plasmid libraries from Step 1 cloning were pre-digested with Esp3I (NEB, R0734L) prior to their use in Golden Gate reactions with the BsmBI-v2 NEBridge® Golden Gate Assembly Kit (NEB, E1602L; note, BsmBI is an isoschizomer of Esp3I) and T4 DNA Ligase Reaction Buffer. Cycling conditions for these reactions were as follows: 42°C x 5 min initial digestion, 65 cycles of 42°C x 5 min digestion followed by 16°C x 5 min ligation, 42°C x 15 min final digestion, 70°C x 20 min heat inactivation, and 10°C hold. Golden Gate reactions were concentrated, Endura Electrocompetent Cells were electroporated, and plasmid libraries were purified as described in the Step 1 library cloning Methods subsection. At this step, each plasmid contains the complete rhodopsin variant expression and barcoded trafficking reporter cassettes, with one barcode-variant combination per plasmid.

### Mammalian cell engineering for Method 2

All cell engineering was performed with a clonal HEK293T cell line constitutively expressing the reverse tetracycline transactivator (rtTA) and containing a Bxb1 recombinase-based landing pad in one copy of the *H11* safe harbor locus. This line was cultured in Gibco DMEM with High Glucose and GlutaMAX Supplement (Thermo Fisher Scientific, 10566024), hereafter DMEM, supplemented with 10% qualified FBS (Thermo Fisher Scientific, 26140079). TrypLE™ Select Enzyme without phenol red (Thermo Fisher Scientific, 12563029) was used for cell dissociation during routine passaging.

For each of the five final plasmid libraries, 5.5e6 cells were seeded into each of ten 10-cm dishes in 12 mL of DMEM + 10% qualified FBS. The following day, cells were co-transfected with 11.8 μg of the appropriate variant plasmid library and 2.92 μg of the Bxb1 recombinase expression plasmid using 29.4 μL of P3000 reagent and 44.2 μL of Lipofectamine 3000 reagent (Thermo Fisher Scientific, L3000075) diluted in Opti-MEM (Thermo Fisher Scientific, 31985088), using incubation times as indicated in the manufacturer’s protocol, for site-specific, single-copy integration of one barcode-variant combination per cell. Three days later, the ten 10-cm dishes were expanded to ten 15-cm dishes without discarding any cells, and, the next day, puromycin (Thermo Scientific, A1113802) was added at 1 μg/mL to each dish to cull cells that did not properly integrate the *RHO* variant expression/reporter cassette. A full media change with fresh 1 μg/mL puromycin was performed three days later. Four days later, cells were dissociated and counted, and 30e6 cells were seeded in 40 mL media into each of four T225 flasks. Three days later, cells were dissociated, counted, and prepared for cryopreservation in BAMBANKER (VWR, 101974-112).

### Assay for Method 2

Individual chunk cell libraries were thawed, allowed to expand, counted, pooled in equal amounts, and plated at 17e6 cells per 15-cm dish, with four replicate dishes per condition. One day later, cells were treated with 1) 2 μM doxycycline hyclate (ApexBio, A4052) to induce rhodopsin variant expression and 2) either DMSO or the YC-001 corrector molecule (MedChemExpress, HY-124717). Approximately 18 hours later, media was aspirated and cells were lysed using 4 mL ice-cold Buffer RLT (Qiagen, 79216) containing beta-mercaptoethanol (Sigma Aldrich, M3148-25ML) for RNase inhibition. Dishes were scraped using cell lifters (Bio Basic, SP91151), lysates were transferred to 5 mL tubes, and lysates were homogenized by repeated aspiration and dispensing using 10 mL syringes with 18-gauge blunt tip needles (VWR, NB18212). Total RNA was purified from 350 μL of the homogenized lysate using the RNeasy Plus Mini Kit (Qiagen, 74134) column-based clean-up that included on-column treatment with DNase (Qiagen, 79254). RNA was eluted in 30 μL of the provided RNase-free water, and concentration and purity were assessed using a spectrophotometer.

cDNA containing reporter gene-encoded barcodes was selectively generated from the purified RNA using a reporter-specific primer and the SuperScript IV First-Strand Synthesis System (Thermo Fisher Scientific, 18091200) according to the manufacturer’s instructions. Reaction products were incubated with RNase A (Thermo Fisher Scientific, EN0531) and RNase H (provided in SuperScript kit) to digest away the original RNA template. NGS libraries were prepared for each sample with PCR using the NEBNext® Ultra™ II Q5® Master Mix and 5 μM forward and reverse primers containing sequences for grafting to an Illumina flow cell (dual indexing). Amplicon sequencing libraries were subjected to 1x26 single-end sequencing using the 50-cycle NextSeq 2000 P3 Reagent kit (Illumina, 20046810) on an Illumina NextSeq 2000 instrument.

**Immunofluorescence staining and flow cytometry of rhodopsin truncating variants** HEK293T cells were maintained in DMEM supplemented with 10% FBS. Prior to the day of transfection,∼1x10^5^ cells were seeded per well of 12-well plate. For transfection, 2 µg of plasmids (WT *RHO*, W126X, G114D, G114X, F116X, F116X-(trunc:348bp) were diluted in Opti-MEM (Thermo) and transfected using Lipofectamine 3000 reagent (Thermo, L3000015) following the manufacturer’s instructions. The following day, cells were dissociated using 0.25% trypsin, diluted and seeded on the chamber slides (Thermo Scientific, 12-565-843) coated with Laminin (Thermo, 23017015, 1:40 dilution in PBS for 1 hour at room temperature). The following day, the cells were washed and fixed with 4% paraformaldehyde (Electron Microscopy Sciences, 15710) and blocked with 3% BSA (Sigma-Aldrich). After blocking, the cells incubated with anti-rhodopsin antibody (Ret-P1 O4886, 1:2000 dilution) for 30 minutes at room temperature. After three washes with PBS, cells were incubated with Alexa Fluor anti-mouse secondary antibody (Thermo, A21121) for 30 minutes at room temperature. After three washes with PBS, the slides were layered and mounted using Prolong glass (Thermo, P36984). The slides were imaged under a fluorescence microscope (Eclipse Ti, Nikon). For flow cytometric analysis, the cells were dissociated and collected 48 hours post transfection, fixed and stained without permeabilization and analyzed using Sony SH900 (see immunostaining and FACS for Method 1).

### Statistical inference of variant effects for Method 2

For bioinformatic and statistical analysis, we applied a broadly similar approach as previously described^63^. Briefly, raw BCL files were demultiplexed with *bcl2fastq*, and barcodes were extracted and counted from the resulting FASTQ files. Raw barcode counts from each sample were joined with the oligonucleotide-barcode map to generate the final count dataset.

To infer variant effects across tested conditions, we applied a negative binomial generalized linear mixed model (NBGLMM). We fit one model per position, where each model included all barcodes derived from variants at that position as well as all wild-type barcodes from the chunk containing that position. The applied model is specified as:

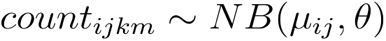

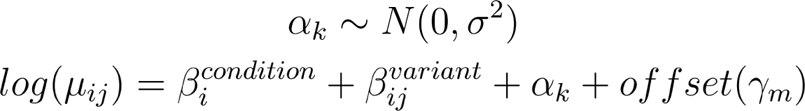

For the *i*th condition, the *j*th variant, the *k*th barcode, and the *m*th sample. In this study, we consider two treatment conditions (DMSO and YC-001) and twenty possible variants at each position. Otherwise, we implement this model as previously described^63^ including fitting with *glmmTMB* and coefficient and marginal mean computation with *emmeans*.

### Additional statistical testing

Statistically significant trafficking score differences were determined using the 95% confidence interval, defined as the mean ± 1.96*(standard error), where the mean and error are returned from the statistical modeling described above. Variant trafficking was significantly lower than wild-type if the upper limit of the 95% confidence interval for variant trafficking was less than 1. Variants were considered to have moderate or high confidence trafficking defects if the upper limit of the 95% confidence interval for variant trafficking was less than 0.7 or 0.5, respectively. Variants were considered to have significant trafficking rescue if the lower limit of the 95% confidence interval for variant trafficking in the presence of 30 μM YC-001 was greater than the upper limit of the 95% confidence interval for variant trafficking at baseline (DMSO vehicle). Conversely, variants were considered to have significant trafficking worsening if the lower limit of the 95% confidence interval for variant trafficking at baseline was greater than the upper limit of the 95% confidence interval for variant trafficking in the presence of 30 μM YC-001.

### Comparison to pathogenicity data and existing computational predictors

To obtain pathogenicity data from ClinVar for known variants, pathogenicity data for all single amino acid missense rhodopsin variants was downloaded February 5, 2025. ClinVar pathogenicity categories were re-coded into three levels: benign, VUS, and pathogenic. The “conflicting classifications” category was excluded. The pathogenicity data were linked to the prediction or trafficking data at the amino acid level. ClinVar ratings with at least “one star” evidence level were used. For computational predictors, pre-computed annotation was obtained for all single-nucleotide changes in the *RHO* gene from the dbNSFP4.9a. Only missense variants were analyzed, which excluded intronic and nonsense variants. After removing 5 sparsely populated predictors, rank scores for 53 predictors were included, ranging from 0 to 1. (Note that while the predictors are named “rankscore”, they are not homogeneously distributed between 0 and 1 like nonparametric ranks would be.) The combined trafficking score scale used in this study (0 = low trafficking, 1 = high trafficking) was reversed and scaled to match the predictors’ rank scores that had a range from 0-1 with 0 as benign and 1 as pathogenic, producing a trafficking_rankscore_homog. The trafficking scores and computational predictor scores were matched to the pathogenicity data at the amino acid level.

## Commercial Relationship Disclosure

Connor H. Ludwig, Nathan Abell, and Matthew L. Albert are current employees of Octant, Inc. Jason Comander is a consultant for Octant, Inc. Kannan V. Manian, Patent application PCT/US2025/010783 (P); Jason Comander, Patent application PCT/US2025/010783 (P).

## Acknowledgments

We acknowledge the Broad Institute of MIT and Harvard, including the Genetic Perturbation Platform (GPP), and Andrew Patentreger and Natan Pirete at the Flow Cytometry Core for library processing and technical advice. The authors thank members of the Ocular Genomics Institute, Massachusetts Eye and Ear for their critical comments, advice, and discussions. We also acknowledge current and former employees of Octant, including Zachary Casey, Henry Chan, Sri Kosuri, Justin Greene, Morgan MacKenzie, Jimin Park, and Maris Kamalu for their technical contributions to the project, as well as Diane Dickel and others at Octant for their critical comments, advice, and discussions.

